# Disrupting central carbon metabolism increases antibiotic susceptibility in *Vibrio cholerae*

**DOI:** 10.1101/2022.12.22.521713

**Authors:** Megan Keller, Xiang Han, Tobias Dörr

## Abstract

Antibiotic tolerance, the ability of bacteria to sustain viability in the presence of typically bactericidal antibiotics for extended time periods, is an understudied contributor to treatment failure. The Gram-negative pathogen *Vibrio cholerae*, the causative agent of cholera disease, becomes highly tolerant to β-lactam antibiotics (penicillin and related compounds) in a process requiring the two-component system VxrAB. VxrAB is induced by exposure to cell wall damage conditions, which results in the differential regulation of >100 genes. While the effectors of VxrAB are relatively well-known, VxrAB environment-sensing and activation mechanisms remain a mystery. Here, we used transposon mutagenesis to screen for mutants that spontaneously upregulate VxrAB signaling. This screen was answered by genes known to be required for proper cell envelope homeostasis, validating the approach. Unexpectedly, we also uncovered a new connection between central carbon metabolism and antibiotic tolerance. Inactivation of *pgi* (*vc0374*, coding for Glucose-6-phosphate isomerase) resulted in an intracellular accumulation of glucose-6-phosphate and fructose-6-phosphate, concomitant with a marked cell envelope defect, resulting in VxrAB induction. Deletion of *pgi* also increased sensitivity to β-lactams and conferred a growth defect on salt-free LB; phenotypes that could be suppressed by deleting sugar uptake systems and by supplementing cell wall precursors in the growth medium. Our data suggest an important connection between central metabolism and cell envelope integrity and highlight a potential new target for developing novel antimicrobial agents.

**Importance:** Antibiotic tolerance (the ability to survive exposure to antibiotics) is a stepping-stone towards antibiotic resistance (the ability to grow in the presence of antibiotics), an increasingly common cause of antibiotic treatment failure. The mechanisms promoting tolerance are poorly understood. Herein, we discovered central carbon metabolism as a key contributor to antibiotic tolerance and resistance. A mutant in a sugar utilization pathway accumulates metabolites that likely shut down the synthesis of cell wall precursors, which weakens the cell wall and thus increases susceptibility to cell wall-active drugs. Our results illuminate the connection between central carbon metabolism and cell wall homeostasis in *V. cholerae* and suggest that interfering with metabolism may be a fruitful future strategy for development of antibiotic adjuvants.

## Introduction

Antibiotic treatment failure is increasingly common in healthcare settings. Antibiotic resistance, the ability of bacteria to grow in the presence of antibiotics, is a major contributor to treatment failure and poses a well-recognized, massive threat to public health. Antibiotic tolerance, which is the prolonged survival of bacteria after antibiotic exposure (1), has also been linked to treatment failure (2) and the development of antibiotic resistance (2–5). Lastly, intrinsic resistance, i.e. the ability to grow in the presence of low concentrations of antibiotic through intrinsic diffusion barriers, target availability, and damage repair functions (sometimes in addition to well-defined resistance factors), likely also contribute to the decreased effectiveness of antibiotics in the clinical setting (6).

While the mechanisms of frank resistance are well-understood, the mechanisms of tolerance and intrinsic resistance remain understudied, preventing us from gaining a more comprehensive insight into the causes of antibiotic treatment failure. Many Gram-negative clinical pathogens are tolerant against antibiotics that target the bacterial cell wall, for example the ordinarily bactericidal β-lactam antibiotics (7). β-lactams inhibit cell wall synthesis, which in susceptible bacteria usually results in degradation of the essential cell wall via the activity of endogenous lytic enzymes, so-called “autolysins” (8). The cell wall consists mostly of peptidoglycan (PG), a complex macromolecule that comprises alternating sugars crosslinked by short peptide chains. This structure normally provides protection against an osmotically variant environment and governs bacterial shape (9). Defying the essentiality of PG, tolerant Gram-negative pathogens survive cell wall degradation induced by antibiotics, and assume a non-replicating spheroplast form that readily reverts to normal growth upon removal of the antibiotic (7, 10–12).

*Vibrio cholerae* is a particularly tolerant enteric pathogen that exhibits high survival upon antibiotic-mediated cell wall degradation, essentially rendering β-lactam antibiotics bacteriostatic agents (11). *V. cholerae*’s tolerance is governed by the VxrAB two-component system (also known as WigKR) (13, 14). Once activated, the inner membrane-localized histidine kinase VxrA phosphorylates the cytosolic VxrB, a transcription factor that alters expression of a large regulon, contributing to the tolerance phenotype. Specifically, upon signal recognition, VxrAB down-regulates genes involved in iron acquisition (15) and cell shape (16), while up-regulating genes involved in peptidoglycan synthesis (15), biofilm formation and motility (17) and Type VI secretion (13). The downstream effectors of this system are well characterized; however, what activates VxrA remains poorly understood. A homologous system in the related bacterium *V. parahaemolyticus* has been proposed to bind β-lactams directly (18, 19), but strong evidence for this is lacking (20, 21). In addition, it is unlikely that β-lactams are the only activators of VxrAB; other structurally distinct antibiotics, overactive cell wall degradation enzymes, and mechanical stress also activate the system (14, 22). With numerous contributors to VxrA induction, we sought to uncover genetic pathways promoting activation of VxrAB to find genes involved in general cell wall homeostasis. In addition to known or expected factors, we found that a disruption in the gene coding for the glycolysis enzyme glucose-6-phosphate isomerase (*vc0374, pgi)* causes strong cell wall defects concomitant with reduced tolerance and intrinsic resistance. Our data showcase a novel connection between glycolysis and cell envelope turnover in the cholera pathogen and open the door for the future development of novel antimicrobial agents that potentiate β-lactam antibiotics by interfering with central carbon metabolism.

## Results

### A screen for induction of the VxrAB regulon identifies expected and novel cell envelope maintenance factors

To identify cell envelope maintenance factors in *V. cholerae* on a genome-wide scale, we screened for transposon-mediated mutational activation of VxrAB using our previously validated, VxrAB-responsive promoters P_vxrAB_-*lacZ* and P_vca0140_-*lacZ* **(Fig. 1A)** on plates containing the chromogenic LacZ substrate X-gal. Putative VxrAB activator Tn insertion mutants (enhanced blue colonies, **Fig. 1A**) were then identified by mapping the transposon insertion sites via arbitrary PCR (see Methods for details). The first round (2 biological replicates of mutagenized cultures) yielded mostly hits in genes with known roles in cell envelope maintenance (*ldcV*, coding for a cytoplasmic L,D carboxypeptidase (23); *vca0040*, a putative undecaprenol pyrophosphate translocase (24); and the main Penicillin-binding protein PBP1a and its activators LpoA and CsiV (25)), internally validating our screen.

**Figure 1.**
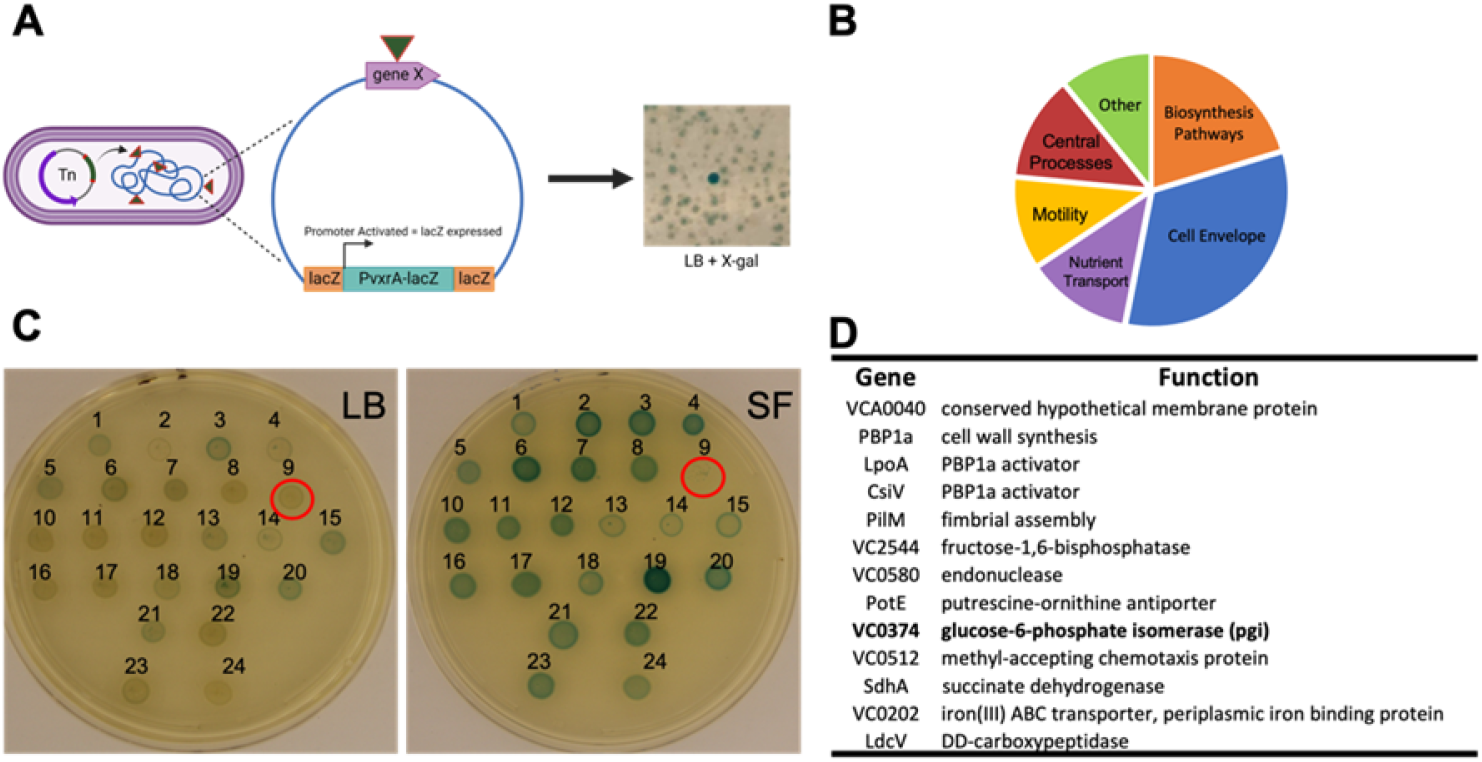
A transposon screen reveals genes required for proper cell envelope homeostasis. A) Schematic depicting the transposon screen strategy B) Pie-chart of the pathways that answered the screen. C) Validation plates of genes of interest on LB and SF plates containing the chromogenic LacZ substrate X-gal. 1= *vc2544#1*, 2= *vca0367*, 3= *pilM*, 4= *toxS*, 5=*vgrR*, 6= *vc0353*, 7=*vc2757*, 8=*vc2683*, 9= *pgi* (indicated by red circle), 10= *vc1911*, 11= *vc2544#2*, 12= *vc0789*, 13= *sdhA#1*, 14= *vc2086*, 15= *sdhA#2*, 16= *fbpB*, 17= *vca0763*, 18= *vc2544*, 19= *vca0137*, 20= *vc2544*#3, 21= *vc2544*#4, 22=*vc0512* 23= P_vxrA_-*lacZ*, 24= P_vca0140_-*lacZ* D) Summary of validated top hits with their assigned functions.

We reasoned that the large number of mutants in PBP1a, LpoA and CsiV might have obscured screen saturation. We thus repeated the screen in the presence of 10 mM D-methionine (which specifically inhibits growth of the PBP1a pathway mutants *pbp1a, lpoA* and *csiV* (26)). This modified screen yielded additional hits, including the gluconeogenesis pathway gene *vc2544* (coding for Fructose 1,6 bisphosphatase) and the glycolysis/gluconeogenesis gene *pgi*, coding for glucose-6-phosphate isomerase (*vc0374*) (**Fig. 1B**). For a full list of genes, see **Table S1**. We validated all hits by plating on X-gal after single-colony purification. Interestingly, we qualitatively noticed that VxrAB background levels appeared elevated in all colonies after conjugational transfer of the transposon shuttle vector. This possibly reflected conjugation-induced cell envelope damage inflicted by the donor strain’s Type IV secretion system (27), resulting in a high number of false positives. To prioritize informative genes further, we thus included growth on salt-free LB (SF), a condition expected to affect the viability of mutants with cell envelope defects, as a validation step. In this additional step, we found that one mutant, a transposon insertion in *vc0374/pgi* had a severe growth defect on both liquid and solid SF, but not standard LB (**Fig. 1C and Fig. S1A**). The transposon mutant in the other glucose metabolism gene that answered the screen, *vc2544* (coding for fructose 1,6, bis-phosphatase, a key step in gluconeogenesis) similarly exhibited enhanced VxrAB induction (**Fig. 1C**), a slight stationary phase morphology defect (**Fig. S1B**) and a subtle growth defect in liquid salt-free LB (**Fig. S1A**); however, the *pgi* defects were more severe. A connection between mutations in glycolysis and cell envelope defects had not previously been reported in *V. cholerae* and we thus decided to focus on the *pgi* mutant for further analysis.

### A pgi mutant exhibits a pronounced cell envelope defect

To validate our transposon screen, we first constructed a clean Δ*pgi* mutant and tested it for growth and cell envelope defects. Consistent with our screen and validation results, Δ*pgi* did not exhibit a growth defect in LB but a severe growth defect in SF, both on plates (**Fig. 2A**) and in liquid medium (**Fig. 2B**). The defect on SF medium could be fully complemented by inducible expression of *pgi*, excluding the possibility of polar effects. To explore if the SF defect was due to reduced osmolarity, or instead specific to the absence of sodium, we plated serial dilutions of a *pgi* mutant culture on SF supplemented with KCl (200mM) or sucrose (180mM). Both adjustments restored growth of the Δ*pgi* mutant (**Fig. 2A**), suggesting that the growth defect is indeed due to low osmolarity. Next, we imaged Δ*pgi* mutant cells grown in LB during stationary phase and exponential phase using phase microscopy. We noted striking morphological defects in the Δ*pgi* mutant, including cell elongation (indicative of a division defect) (**Fig. 2C**); these defects were more pronounced during rapid exponential growth and partially resolved during stationary phase. Cell area and length were calculated using ImageJ and significantly differed between WT and the Δ*pgi* mutant (**Fig. 2D**). These results support the idea that the Δ*pgi* mutation causes defects in cell envelope homeostasis.

**Figure 2.**
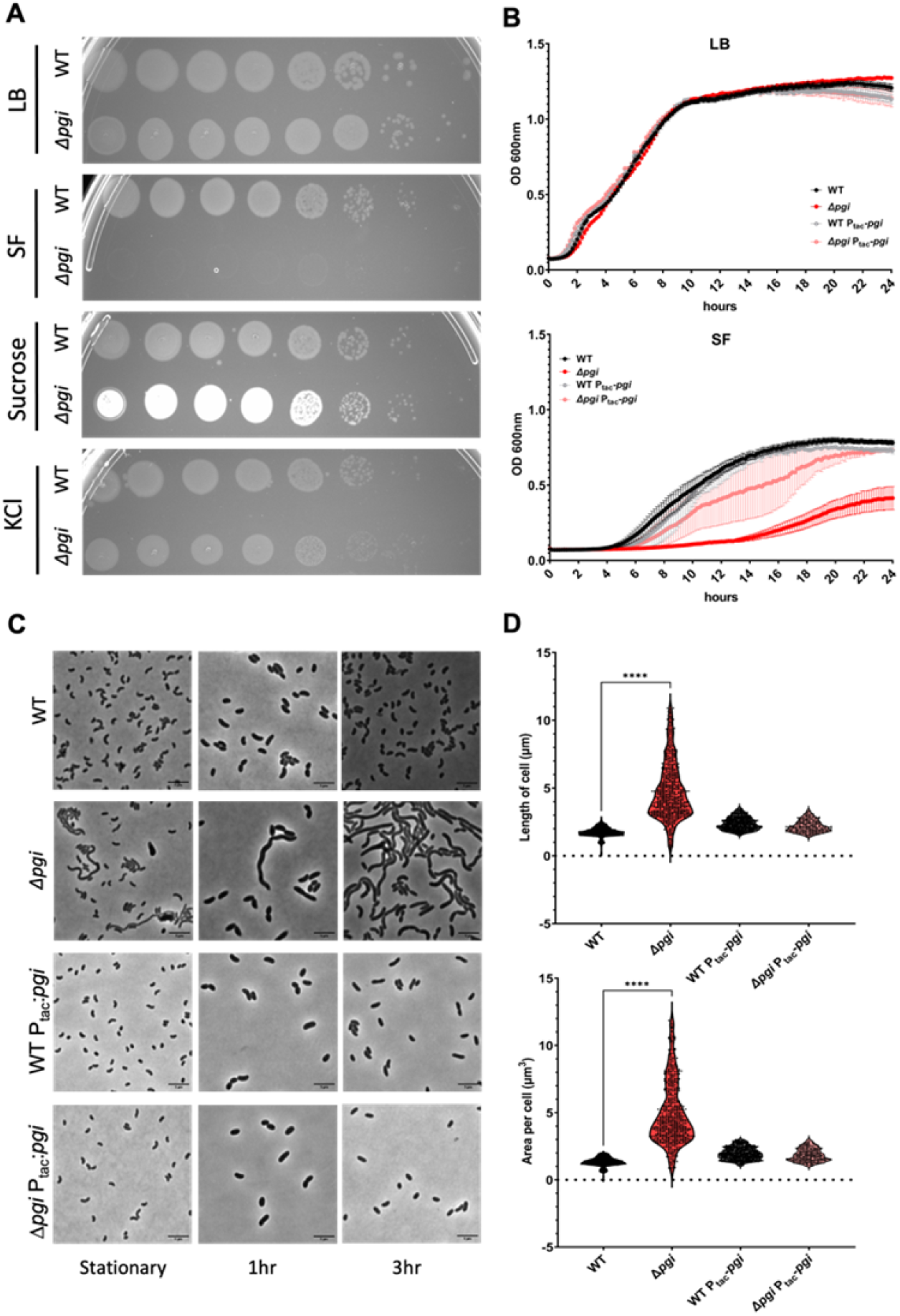
A *pgi* mutant exhibits severe cell envelope defects during growth in LB. A) Overnight cultures of the indicated strains were serially diluted and spotted on LB, SF, SF + 180mM sucrose, and SF + 200mM KCl. B) Growth curves of WT and *Δpgi* in LB or SF liquid media. The indicated strains were diluted 1:100 from overnight cultures and analyzed on a growth curve analyzer (BioScreen). Data represent mean +/- standard error (n=3 independent biological replicates). C) The indicated strains were diluted 100fold into fresh LB, then imaged on an agarose pad after 1 and 3 hours of growth. Scale bar = 5μm D) Quantification of cell area and width conducted with MicrobeJ, n > 250 individual cells.

### The pgi mutant exhibits reduced tolerance and resistance to cell wall-acting antibiotics

We reasoned that the *pgi* mutant’s cell wall defect might enhance susceptibility to cell wall-acting antibiotics; indeed, *pgi* previously answered our TnSeq screen for tolerance defects in *V. cholerae* (10). We next performed a zone of Inhibition (ZOI) assay using a panel of antibiotics, to further parse this out. Consistent with its cell wall defect, the Δ*pgi* mutant was significantly (p<0.001 using Welch’s T-test) more sensitive to β-lactam antibiotics such as PenG, meropenem, and mecillinam, while retaining little to no sensitivity differences to antimicrobials with other mechanisms of action, i.e. hydrogen peroxide, colistin, and tetracycline. (**Fig. 3A**). Susceptibility to antibiotics that are typically excluded from Gram-negative bacteria by the outer membrane permeability barrier (vancomycin and ramoplanin) was also slightly, but significantly increased in the *pgi* mutant, suggesting increased outer membrane damage.

**Figure 3.**
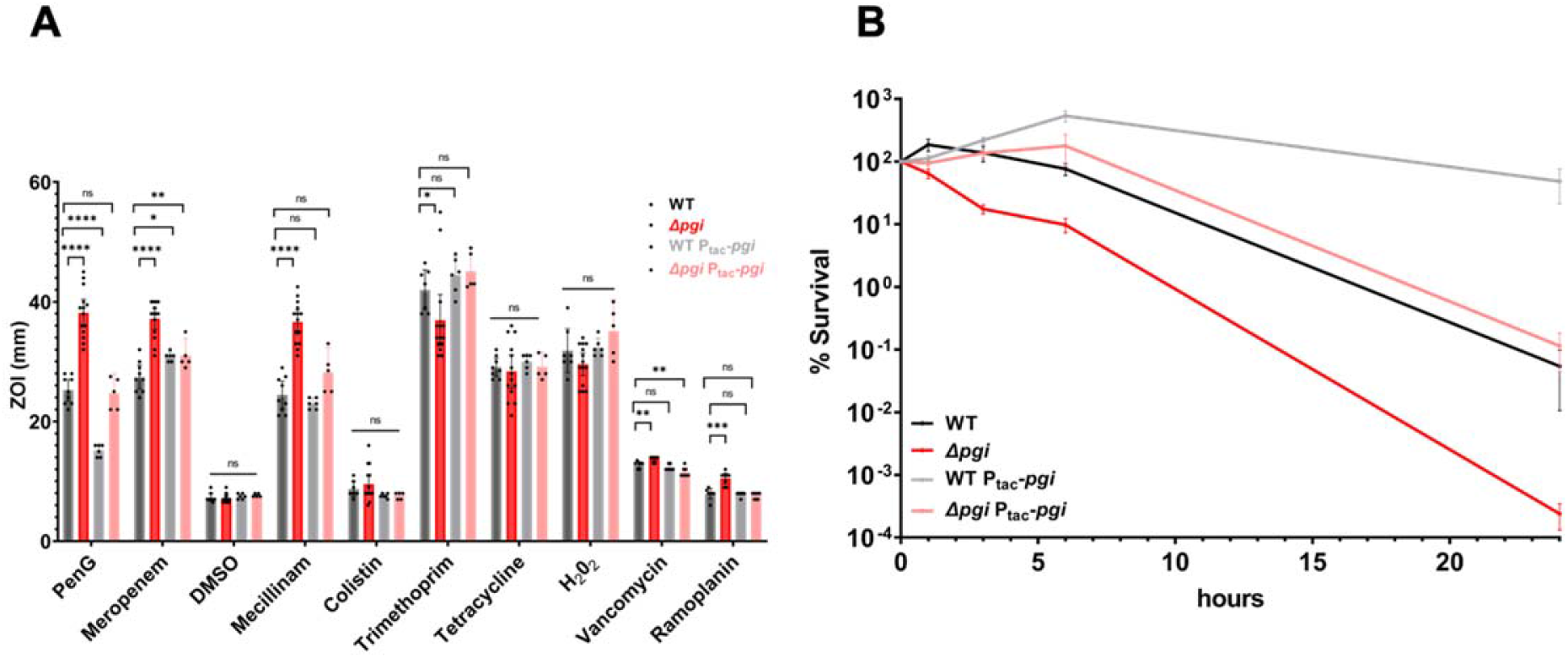
A *pgi* mutant exhibits decreased tolerance and resistance to cell wall-active antibiotics. A) Zone of inhibition measurements from a disk diffusion experiment. Data represent at least 5 independent biological replicates; raw data points are shown with bars indicating 95% confidence interval. Statistical significance was assessed via Welch’s T-test **** = p <0.0001. B) Time-dependent killing experiment. Bacteria were grown to exponential phase and then treated with 100 μg/mL PenG (10x MIC). At the indicated timepoints, serial dilutions were plated on LB and grown overnight at 30 °C. Data represent the means of 6 independent biological replicates +/- standard error.

To next test the *pgi* mutant’s effect on antibiotic tolerance, we conducted a time-dependent killing experiment using PenG (100 μg/mL, 10 x MIC). While viability of the WT strain decreased only slightly during the experiment, consistent with our previous observations (10), Δ*pgi* exhibited a dramatic, 10,000-fold drop in survival over the course of 24 hours (**Fig. 3B**). Interestingly, overexpression of *pgi* increased survival in the WT and over-complemented the *pgi* mutant. Similarly, *pgi* overexpression increased resistance against penicillin (but not other cell wall-active antibiotics) in the WT. These results suggest that *pgi* levels are limiting for both optimal antibiotic tolerance and resistance to penicillin even in the wild type. Collectively, our data suggest that *pgi* plays a central role in intrinsic resistance and tolerance against cell wall-acting antibiotics.

### Suppressor mutations in sugar transport system components restore pgi defects

While conducting the previously mentioned plating and kinetic experiments in SF medium, we noticed a high frequency of spontaneous suppressor mutants arising in the *Δpgi* background. With the goal of gaining more mechanistic insight into *pgi’s* role in cell wall homeostasis, we purified these mutants to study their effects. Whole genome sequencing revealed a similar set of suppressors in 3 independent rounds of selection. Of the 67 suppressors we isolated and whole-genome-sequenced, 52 mapped to the *ptsI* gene (*vc0965*), coding for the first protein in the phosphate relay system that ultimately effects sugar phosphorylation during phosphotransferase-mediated uptake. The remaining suppressors mapped to another component of the PTS system, *vc0964* (crr, coding for the EIIA component), *treB*, encoding a putative trehalose import component, and two genes involved in maltose transport, *vca0944* and *vca0011* (**Table S2**) (28). Our initial validation revealed that these mutants at least partially suppressed all phenotypes associated with the *pgi* mutation, i.e. the SF growth defect, morphological defects upon culturing in LB, as well as antibiotic sensitivity phenotypes, despite selection in SF only (**Fig. S2A-C**).

The identity of suppressor mutations implied likely loss-of-function. To validate suppression of *Δpgi* phenotypes, we constructed clean deletions and complementation strains in the WT and Δ*pgi* backgrounds, focusing on *ptsI* and *treB* **(Fig. 4A)**. These mutations indeed restored viability of the Δ*pgi* strain to varying degrees. Deletion mutants in *ptsI* completely rescued the Δ*pgi* SF sensitivity phenotype and morphological defects (**Fig.4B-C**), while Δ*treB* promoted partial restoration of morphology and cell viability, with complete restoration of intrinsic resistance as measured by ZOI assay (**Fig.4B-C**). Consistent with previous work in *E. coli* and *S. enterica* (29), our suppressor analysis thus overwhelmingly suggests that the *pgi* mutant suffers from sugar phosphate toxicity, which can be mitigated by curbing sugar uptake.

**Figure 4.**
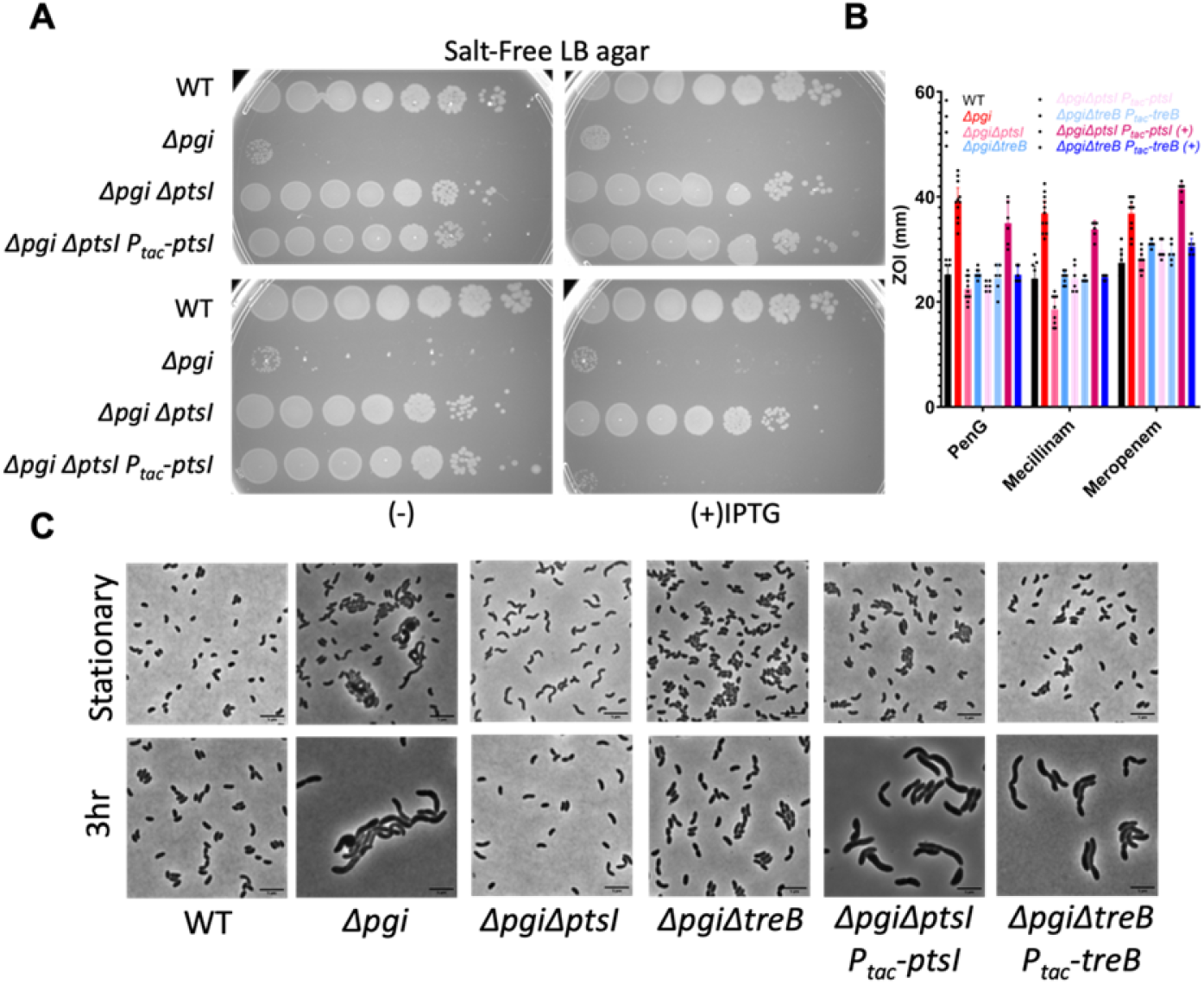
Mutants in sugar uptake systems suppress Δ*pgi* cell envelope defects. A) The indicated mutant strains carrying IPTG-inducible complementation constructs were serially diluted and plated on SF agar with or without IPTG. B) Zone of inhibition measurements from a disk diffusion experiment. Data represent 6 independent biological replicates; raw data points are shown. Statistical significance was assessed via Welch’s T-test **** = p <0.0001. C) The indicated mutant strains carrying IPTG-inducible complementation constructs were imaged after growth in exponential phase in LB with (+) inducer.

### Untargeted metabolomics reveal accumulation of glycolysis intermediates in the *pgi* mutant

We next sought to determine the specific compound that results in Δ*pgi* defects. To examine the metabolite differences in the *pgi* mutant versus WT, we conducted untargeted metabolomics of both strains during their exponential growth in LB, where the mutant exhibited the most drastic phenotype. Using both positive and negative Hydrophilic Interaction Liquid Chromatography (HILIC) to probe both central metabolism and cell envelope precursors based on their surface charges, over 800 metabolites appeared altered between WT and the Δ*pgi* mutant **(Fig. S3)**. Multivariate Analysis was done and the PCA (Principal component Analysis) showed a clear differentiation between WT vs. Δ*pgi* groups. Glycolysis products like β-D-Fructose-6-phosphate (6-o-phosphono hex-2-xylofuranose) and glucose-6-phosphate were dramatically (53- and 236-fold, respectively) increased in the *pgi* mutant. The metabolic intermediate phloroglucinol was also highly (>100fold) increased, though the significance of this observation is unclear (*V. cholerae* lacks a homolog of the PhlD synthetase of the only bacterium known to produce phloroglucinol, *Pseudomonas fluorescens* (30)). These results are partially consistent with *pgi* mutants in *E. coli* and other Gram-negative species grown in the presence of glucose, where an accumulation of G6P was likewise observed (29). The accumulation of both G6P and F6P suggests that both glucose import (indicated by the accumulation of G6P) and gluconeogenesis (accumulation of F6P) are active during growth in LB medium. Alternatively to gluconeogenesis, the F6P accumulation could be due to enhanced flux into the pentose phosphate pathway (PPP) (similar to the *E. coli pgi* mutant, (31)). While metabolomics did not reveal any typical PPP intermediates, enhanced metabolic flux into PPP can in principle result in the generation of reactive oxygen species (ROS) (31), which in turn might affect Δ*pgi*’s viability on salt-free LB. To test this, we plated the *pgi* mutant on SF plates containing catalase (which restored growth to a Δ*oyxR* Δ*katB* Δ*katG* hypersensitive mutant as a positive control for effective catalase activity in our plate assay, **Fig. S4**). Catalase did not restore growth to Δ*pgi*, suggesting that ROS do not contribute to the mutant’s cell wall defect.

The accumulation of G6P was surprising, as LB medium does not contain any added glucose and has a very low fermentable sugar content (32). To test if LB contains residual glucose, we used a glucose quantification kit to assess glucose levels in our LB medium. Surprisingly, we found that this medium does contain significant amounts of glucose (3 μg/mL (**Fig. S5B**)), likely explaining the *pgi* mutant’s phenotypes. However, our suppressor screen was also answered by *treB*, coding for a component of the trehalose PTS uptake system. This could either indicate TreB’s moonlighting ability to also import glucose, or suggest the presence of trehalose in LB. Trehalose can be imported by *V. cholerae* and converted to glucose-6-phosphate by the hydrolase TreC (33). Residual yeast-derived trehalose in LB could in principle contribute to glucose accumulation in the *pgi* mutant, and this would be consistent with TreB answering our *pgi* SF suppressor screen (**Fig. 4**). To explore this possibility further, we first plated the *pgi* mutant on MM and SF plates containing trehalose. Similar to glucose plates, Δ*pgi* failed to grow on trehalose and this could be partially suppressed by deletion of *treB*, raising the possibility that trehalose might contribute to *Δpgi* mutant phenotypes **(Fig. S5A)**. However, direct quantification of trehalose using a commercial kit failed to reveal significant trehalose in LB, suggesting either concentrations below the limit of detection (0.1-8.0 μg/L), or indeed a lack of trehalose.

### Glucose toxicity in the *pgi* mutant may contribute to reduced cell wall precursor synthesis

Next, we asked how the accumulation of these glycolysis metabolites may cause the observed cell wall defects. Glucose-6-phosphate accumulation is associated with cell wall stress in other bacteria and even fungi (34, 35). G6P can be converted to G1P by *pgcA* (also called *pgm* in E.coli); in *Bacillus subtilis*, G1P likely inhibits the cell wall precursor synthesis enzyme, GlmM (36), which converts glucosamine-6-phosphate (Gln6P) to glucosamine-1-phosphate (Gln1P), an important step in the synthesis of the essential PG precursor UDP-N-acetylglucosamine. We hypothesized that a similar GlmM poisoning by G1P might at least partially explain the *pgi* mutant’s cell wall defects. According to our metabolomic data, the *pgi* mutant accumulates (albeit slightly) Gln6P, the natural substrate for GlmM, perhaps indeed suggesting reduced GlmM activity. We reasoned those high concentrations of G6P (or, as in *B. subtilis*, Glucose-1-phosphate) might competitively inhibit GlmM activity (or another precursor synthesis enzyme) due to its structural similarity to the natural substrate Gln6P. If this was the case, externally supplying Gln6P might relieve this inhibition. *V. cholerae* lacks a direct importer for glucosamine but can convert external GlcNAc to Gln6P (28). We thus supplied the Δ*pgi* mutant with GlcNAc, which in *V. cholerae* gets imported via the NagE transporter and then converted into Gln6P via NagA (37, 38). Addition of GlcNAc to either SF LB or M9 glucose minimal medium (where the *pgi* mutant exhibited a drastic growth defect) indeed fully rescued growth (**Fig. 5**), which is in line with an inherent reduction of PG precursor synthesis being at the heart of *pgi* mutant phenotypes.

**Figure 5.**
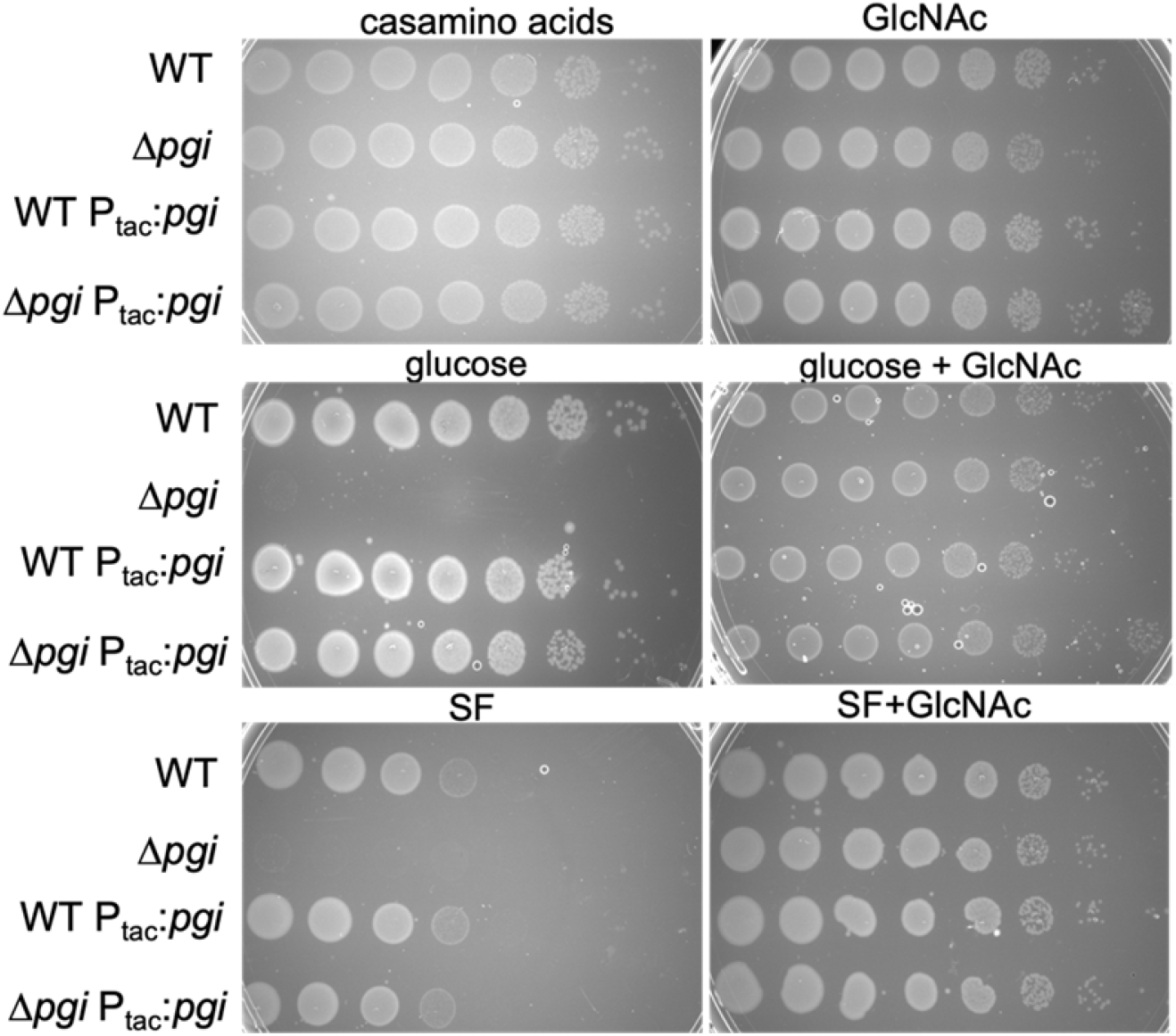
Supplementation with N-acetyl glucosamine suppresses Δ*pgi* mutant phenotypes. Serial dilutions of the indicated strains were plated on salt-free LB (SF) or M9 minimal media supplemented with casamino acids (0.2 %) or glucose (0.2 %), with or without addition of 0.2 % N-acetyl glucosamine (GlcNAc).

## Discussion

Herein, we present data demonstrating that disrupting glycolysis interferes with optimal cell wall homeostasis in *Vibrio cholerae* (**Fig. 6**). Deletion of a key enzyme in glycolysis and gluconeogenesis, *pgi*, caused severe morphological defects and increased sensitivity to low osmolarity and cell wall targeting antibiotics. Suppressors of this defect mapped to sugar uptake systems, with partial suppression afforded by inactivation of the trehalose uptake system. In addition, we found that the *pgi* mutant accumulates G6P and F6P during growth in LB, with indirect evidence that minimal G6P might also be derived from trehalose present in LB. While LB is generally considered a medium virtually devoid of sugars (and especially glucose) (32), our data demonstrate that there is in fact glucose in LB and at least *Vibrio cholerae* might potentially convert other components of this rich medium (e.g., trehalose) to glucose – *caveat experimentator!* (32). Interestingly, a recent study demonstrated that a *ptsI* mutation reduces the length of *E. coli* cells grown in LB broth, likewise suggesting a role for sugar uptake in cell division and/or elongation (39). These data seem to be in line with what we report here – both because they suggest that some glucose is available in LB broth, and because they imply a connection between sugar metabolism and cell envelope homeostasis and cell growth.

**Figure 6.**
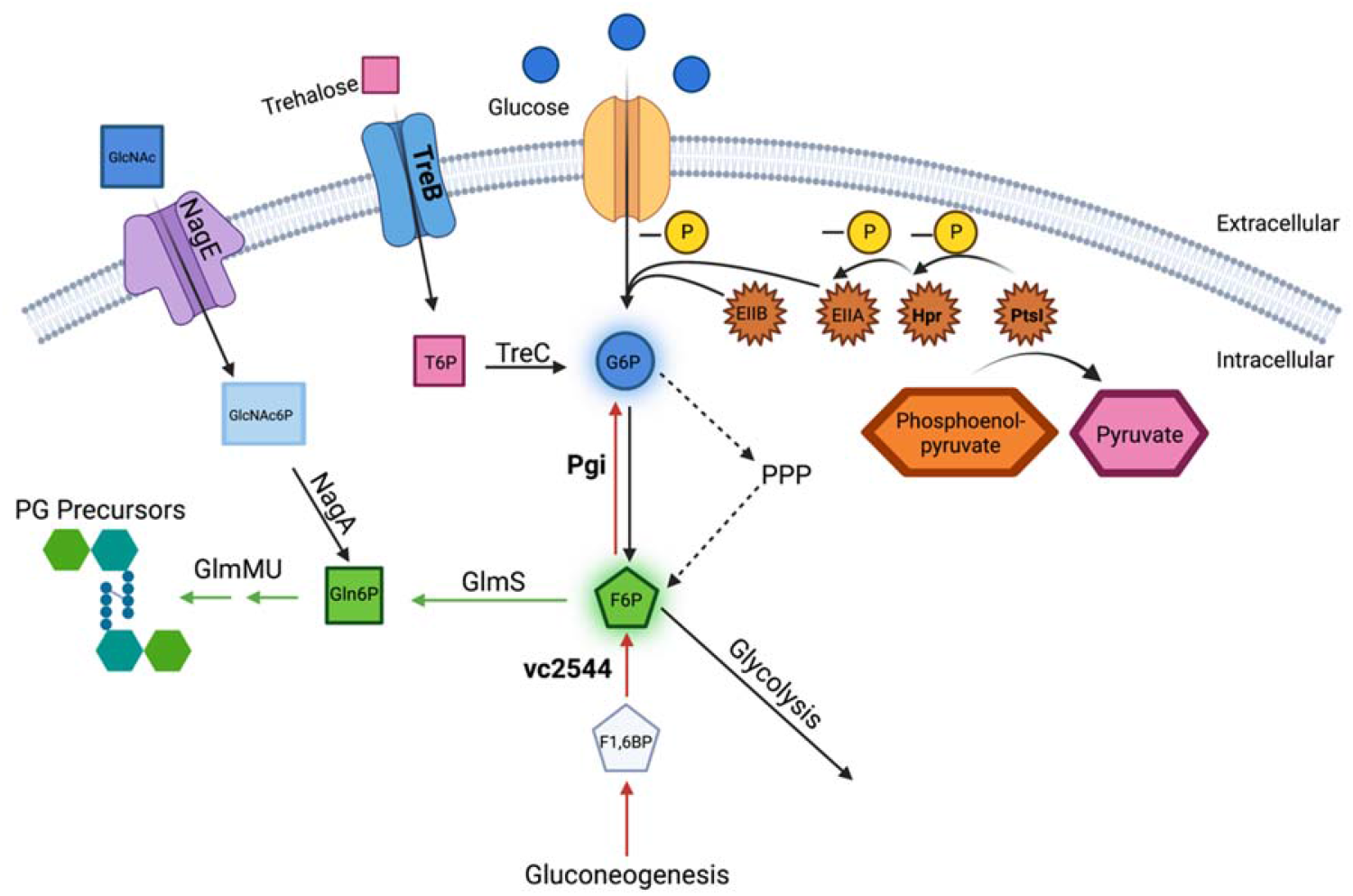
Overview model of the coordination between central metabolism and PG precursor synthesis. Extracellular glucose and trehalose can be up taken into the cell through PtsG and TreB, respectively. Trehalose is converted to Trehalose-6P (T6P) by TreC. Glucose is phosphorylated via the PtsI/Hpr/EIIA(crr)/EIIB phosphorelay cascade, which is initiated by the phosphate transfer from PEP to PtsI. Pgi converts G6P to F6P during glycolysis (black) and vice versa during gluconeogenesis (red). The Pentose Phosphate Pathway (PPP) also contributes to the generation of F6P from G6P. F6P can be converted to Glucosamine-6P (Gln6P) by GlmS and then further to cell wall precursors by GlmMU. Extracellular GlcNAc can be up taken and phosphorylated by NagE and the PTS system to GlcNAc-6P. This is then converted to Gln6P by NagA. Genes answering the screen for VxrAB are bolded and metabolites that increased in the *pgi* mutant are highlighted.

Our data shed new light on the emerging connection between central metabolism and cell wall homeostasis. Mutants disrupted in carbon flux have been extensively characterized in model organisms like *E. coli* and *B. subtilis*. In *B. subtilis*, for example, a checkpoint protein, *cpgA*, is impacted by glucose toxicity. When deleted, a cascading imbalance of metabolites contributes to accumulation of 6-phosphogluconate and subsequent shutdown of both pentose phosphate pathway and glycolysis by simultaneous inhibition of 6-phosphogluconate dehydrogenase (GndA) and *pgi*, resulting, among other defects, in cell envelope damage (40). Also in *B. subtilis*, mutating *pgi* and glucose-6-phosphate dehydrogenase (i.e. this mutant is defective in both glycolysis and PPP) results in the accumulation of Glucose-1-phosphate, which causes apparent cell wall synthesis inhibition and ultimately cell lysis (36). This was shown to be the consequence of conversion of G6P (accumulating in the double mutant) to G1P via the PgcA enzyme; G1P subsequently inhibits an unknown step in PG precursor synthesis, likely GlmM (36). While our data do not reveal the exact mechanism of cell wall perturbations in the *pgi* mutant, a similar metabolic poisoning of GlmM or another precursor synthesis enzyme might be the culprit. In principle, cell wall precursor synthesis might also be reduced in the *pgi* mutant due to its diminished ability to generate F6P, the branching point between glycolysis and PG precursor synthesis. However, our metabolomics revealed a substantial accumulation of F6P in the *pgi* mutant (53-fold compared to WT), rather than a decrease, suggesting that either PPP or gluconeogenesis can efficiently generate F6P, and/or that the effective shuttling of F6P into PG precursor synthesis is blocked at the level of GlmS activity. Interestingly, a *pgi* mutant has recently also been implicated in suppression of arabinose-toxicity in *V. cholerae* (41). At high concentrations, arabinose induces cell wall damage in an apparently *pgi*-dependent way. While the mechanistic details of this observation are unclear, this likewise implies a connection between central carbon metabolism and cell wall homeostasis. Deciphering the exact mechanism of cell wall perturbations in the *pgi* mutant will be part of a future study. Interestingly, our original screen was also answered by a mutant in the gluconeogenesis enzyme fructose 1,6 bisphosphatase. This mutant had a more subtle phenotype than the *pgi* mutant, perhaps indicating that disruption of gluconeogenesis causes a decrease in F6P levels (which is only partially buffered by glycolysis using LB’s limited glucose, or by F6P generation through the pentose phosphate pathway), and the associated decrease in PG precursor availability might be the underlying cause of this mutant’s cell wall defects.

Consistent with internal inhibition of cell wall synthesis, our data also demonstrate that G6P accumulation promotes increased susceptibility to cell wall acting antibiotics, reducing both intrinsic resistance and tolerance to cell wall-acting agents in the cholera pathogen. Our data thus add support to the emerging model that interfering with central metabolism is a promising strategy to potentiate antibiotics (42). Crucially, glycolysis flux as a modulator of cell wall homeostasis appears to be conserved from bacteria to fungi (34, 35), perhaps suggesting that this represents an ancient homeostatic feedback mechanism between nutrient state of the cell and cell envelope integrity that could be exploited as a broad target for the development of antimicrobials.

## Methods

### Bacterial Strains and Growth Conditions

All *V. cholerae* strains used in this study are derivatives of *V. cholerae* El Tor strain N16961 listed in Appendix Table S1. *V. cholerae* was grown on Luria-Bertani (LB) medium, in Salt-Free LB (for 1L bottle, 10g Tryptone, 5g yeast extract, 10g NaCl, with addition of 15g agar for solid medium), or in M9 Minimal medium with 0.1% Casamino acids at 30°C unless otherwise indicated; 200ug/mL of streptomycin was also added (N16961 is streptomycin resistant). Where applicable, growth media were supplemented with 0.2% glucose, 0.2% trehalose, or 0.2% GlcNAc. For growth experiments, overnight cultures were diluted 500fold into 1mL LB + streptomycin and incubated in 100well honeycomb wells in a Bioscreen growth plate reader (Growth Curves America) at 37°C with random shaking at maximum amplitude, and OD_600_ recorded at 10 min intervals.

### Plasmid and Strain Construction

Oligos used in this study are summarized in Appendix Table S1. *E. coli* DH5α λ*pir* was used for general cloning, while *E. coli* MFDλ*pir* (a diamino pimelic acid auxotroph) or SM10 λ*pir* were used for conjugation into *V. cholerae* (43). Plasmids were constructed using Gibson assembly (44). All plasmids were Sanger sequence verified. The transcriptional fusion plasmid pAW61 was used to create the P_vxrAB_-lacZ and P_vca0140_-lacZ reporter strains. To this end, 500bp regions upstream of the genes *vca0565* (vxrA) or *vca0140* were amplified from N16961 genomic DNA by PCR and cloned into the pAW61 vector, a suicide plasmid containing *E. coli lacZ* downstream of a multiple cloning site, flanked by *V. cholerae* lacZ upstream and downstream sequences, as well as the *sacB* marker for counterselection. pAW61 was then used to deliver the reporter construct into the *V. cholerae* chromosome, replacing native lacZ with the promoter:*lacZ* fusion. Conjugation of pAW61 into *V. cholerae* was performed by mixing overnight cultures 1:1 (100 μL donor + 100 μL recipient) in 800 μL fresh LB, followed by pelleting (7000 x rpm, 2 minutes) and resuspending in 100 μL LB. The mixture was then spotted onto LB agar (+ 600 μM diamino-pimelic acid, DAP, for E.coli growth) and incubated for 4 hours (overnight for pTOX5 deletions) at 37 °C. Selection for single crossover mutants was then achieved by streaking the mating mixture on streptomycin (200 μg/mL) +carbenicillin (100 μg/mL) in the absence of DAP. Deletion mutants were then obtained by counterselection on salt-free LB + 10 % sucrose (to inhibit growth of *sacB*+ single crossover mutants). Mutants were then verified by colony PCR using primers 41 (AIW567) and 8 (MKchromolacZ rev) (Appendix Table S2).

Gene deletions were constructed using the pTOX5 *cmR/msqR* allelic exchange system (45). In short, 500 bp regions flanking the gene to be deleted were amplified from N16961 genomic DNA by PCR, cloned into suicide vector pTOX5, and conjugated into *V. cholerae*. The first round of selection after mating on LB + DAP + 1 % glucose was performed on LB + chloramphenicol (5 μg/μg mL) + streptomycin + 1% glucose at 30°C. Chloramphenicol-resistant colonies were picked into Eppendorf tubes with 1mL LB + 1% glucose and incubated at 37°C without agitation for 3hr, then pelleted and counter-selected on M9 + 2% rhamnose at 30°C. Deletions were verified by PCR. Since the *pgi* mutation confers glucose sensitivity, we created mutants in this background without glucose supplementation and then verified plasmid loss with patch plating (+)/(-) chloramphenicol. Successful knockouts were then verified using flanking and internal primers respectively and verified with whole genome sequencing.

Complementation strains were created using the chromosomal integration plasmid pTD101, a derivative of pJL1 (46) containing lacI^q^ and a multiple cloning site under control of the IPTG-inducible P_tac_ promoter. pTD101 integrates into the native *V. cholerae lacZ* (*vc2338*) locus. Genes for complementation experiments were amplified from N16961 genomic DNA, introducing a strong consensus RBS (AGGAGA), and cloned into pTD101 using Gibson assembly. pTD101 was integrated into the *V. cholerae* chromosome as described above for pAW61 and colony PCR verified using primers 5 and 8.

### Transposon Mutagenesis

Overnight cultures of P_vxrAB_-lacZ, P_vca0140_-lacZ, and *E. coli* SM10 carrying the mariner transposon delivery plasmid pSC189 (47), were washed, conjugated in equal volumes and spotted onto 45 μm filter discs placed on LB plates. These were incubated for 4hr at 37°C, resuspended in 2mL of LB, plated onto large LB + Xgal + kan (50ug/mL) + streptomycin (200ug/mL) plates, and grown overnight at 37°C. Blue colonies (“vxrAB on”) were picked and purified on the same medium overnight at 30°C. Single colonies were then picked and resuspended in a 96-well plate with 100uL LB + streptomycin. Arbitrary PCR was conducted using ARB 1, ARB 2, Himarout, and H1 primers (Table S1). The first PCR utilized ARB1 and Himarout, adding external oligos to the transposon specific sequences. 1.5μl of PCR product was added to PCR 2, where H1 and Arb2 were used to make internal complementary sequences. Transposon-genomic DNA junctions were identified using Sanger sequencing with H1 primers followed by nucleotide BLAST.

### Cell Viability Assay

To test cell viability, overnight cultures were added to sterile 1X PBS for serial dilution from 1:10 to 1:10^7^. 5uL of overnight cultures and diluted cultures were spotted for CFU/mL on different media plates, as described in the figure legend. Dried plates were then incubated at 30°C overnight and counted the next day. For time-dependent killing experiments, overnight cultures of wild-type and mutant strains were diluted 1:100 and grown in LB medium at 37°C for 1.5hr and then supplemented with 100μg/mL PenG and continued to grow at 37°. At designated time points, samples were collected, serially diluted using sterile 1xPBS, and plated for CFU/ml on LB plates. Plates were then incubated at 30°C overnight and counted the next day.

### Antibiotic Sensitivity Assay

For zone of inhibition assays, a lawn of overnight cultures (100 μL) was spread on an LB plate and allowed to dry for 15 min. 10 μL of antibiotic solutions (100 mg/mL PenG; 10 mg/mL meropenem; 20 mg/mL mecillinam; 12 mg/mL colistin; 100% DMSO; 50 mg/mL trimethoprim; 5 mg/mL tetracycline; 100% hydrogen peroxide; 100 mg/mL vancomycin; 100 mg/mL ramoplanin) were placed on filter disks (6 mm) onto the agar surface and incubated at 37°C overnight before measurements. Statistical analysis was performed using Welch’s T-test.

### Microscopy

Strains were grown as previously described and imaged without fixation on LB 0.8% agarose using a Leica DMi8 inverted microscope. Phase-contrast images were analyzed using MicrobeJ, an ImageJ plug-in. Default parameter settings were applied.

### Spontaneous Suppressor Identification

Spontaneous suppressors were obtained from the *pgi* mutant on salt-free LB, followed by incubation at 37 °C overnight. Colonies were single-colony purified, grown overnight in LB and then validated for stability of the mutation by renewed growth in salt-free LB. Validated suppressors we identified using whole-genome sequencing (Microbial Genome Sequencing Center (MiGS, Pittsburg, PA) using NCBI: NC_002505 NC_002506 as the reference genomes.

### Metabolomics

Samples were diluted 1:100 from stationary growth into LB and grown for 1.5 hrs at 37°C to reach exponential growth. They were centrifuged at 750 x g for 5 min and resuspended in 80% methanol. They were kept at −80 °C and transferred to the Metabolomics Core (Cornell University). There, the samples were centrifuged at 14,000 x g for 10 min at 4–8°C to pellet the cell debris. Then, the metabolite-containing supernatant was transferred to a new 15-mL conical tube on dry ice and lyophilized to a pellet using no heat. Compound Discoverer was used for data analysis of untargeted metabolomics in positive and negative mode, comparing the m/z in the samples with different databases: Chemspider, bioCyc, HMDB, Food metabolome, massbank, Lipidmaps, Mzcloud. 425 metabolites were annotated using the negative HILIC and filtered after normalization and removing the background and false positives. 328 annotated metabolites were filtered based on MS2 spectra, peak rating and with +/- 5 ppm error. 378 metabolites were annotated using the positive HILIC and filtered after normalization and removing the background and false positives, 341 annotated metabolites were filtered based on MS2 spectra, peak rating and with +/- 5 ppm error. Multivariate Analysis was done and the PCA (Principal component Analysis) showed a clear differentiation between both groups. QC (pooled samples) was used for normalization across all samples and QC (group) samples were used for identification.

### Quantification of sugars

Glucose in LB was quantified using the Glucose (GO) Assay Kit from Sigma-Aldrich (Catalog Number GAGO20) following manufacturer’s instructions. In summary, D-Glucose is converted into D-gluconic acid and hydrogen peroxide. The basic hydrogen peroxide is then mixed with reduced o-dianisidine and a colorimetric reaction occurs that is measured at 540nm. A standard curve was created using known glucose concentrations. Trehalose was measured using the Meganzyme kit K-TREH following manufactures instructions.

## Acknowledgements

Tolerance research in the Dörr lab is supported by NIH R01-AI143704. MK is supported by NSF GRFP #DGE-1650441. We thank John Helmann and Yesha Patel for critical comments on the manuscript.

## Supplementary Figure and Table Legends

**Supplemental Figure 1:**
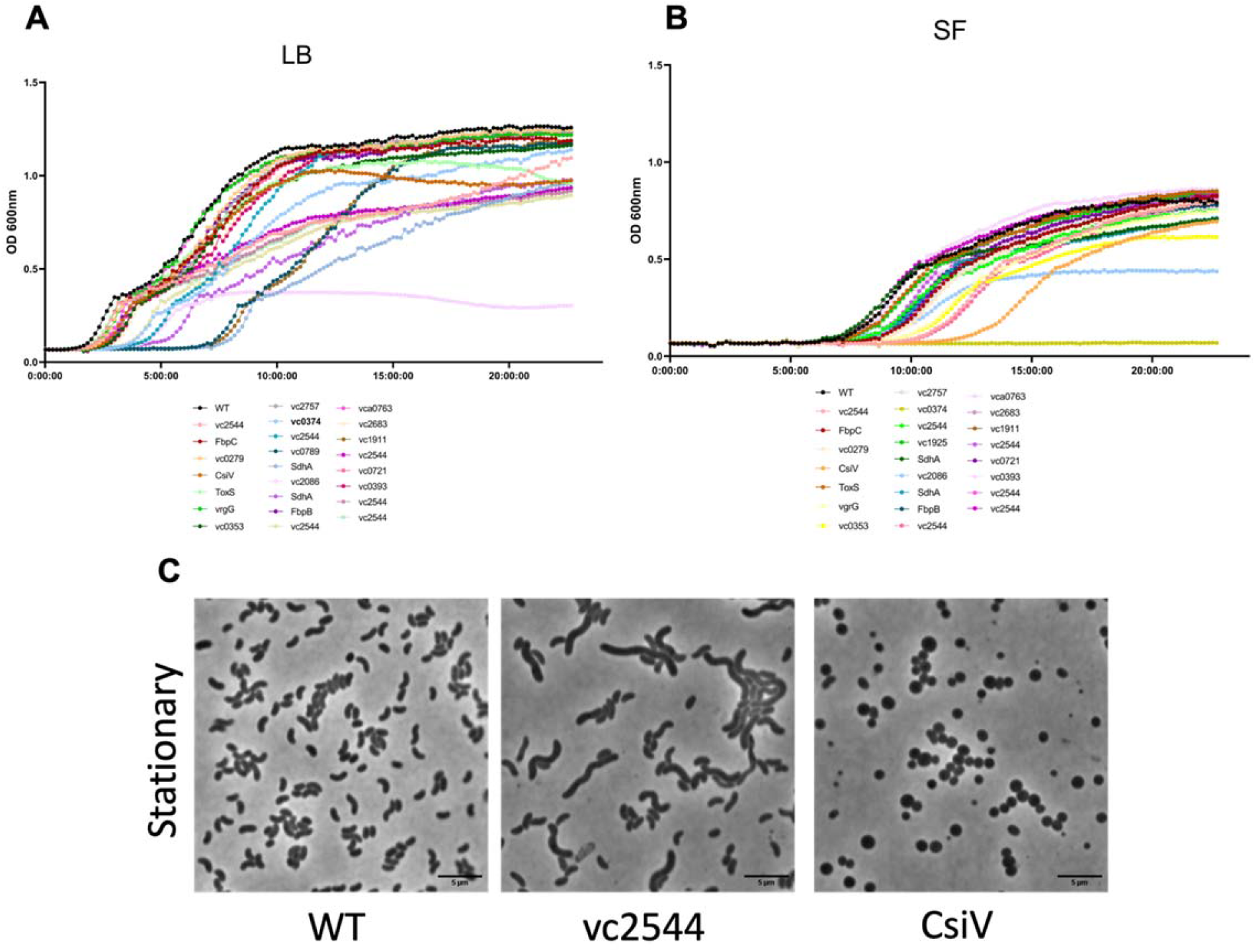
Preliminary validation of transposon hits by growth dynamics and morphology. A) Growth curves with either LB or SF media of 24 transposon hits compared to WT. B) Microscopy of overnight cultures of selected mutants compared to WT. Scale bar = 5μm.

**Supplemental Figure 2:**
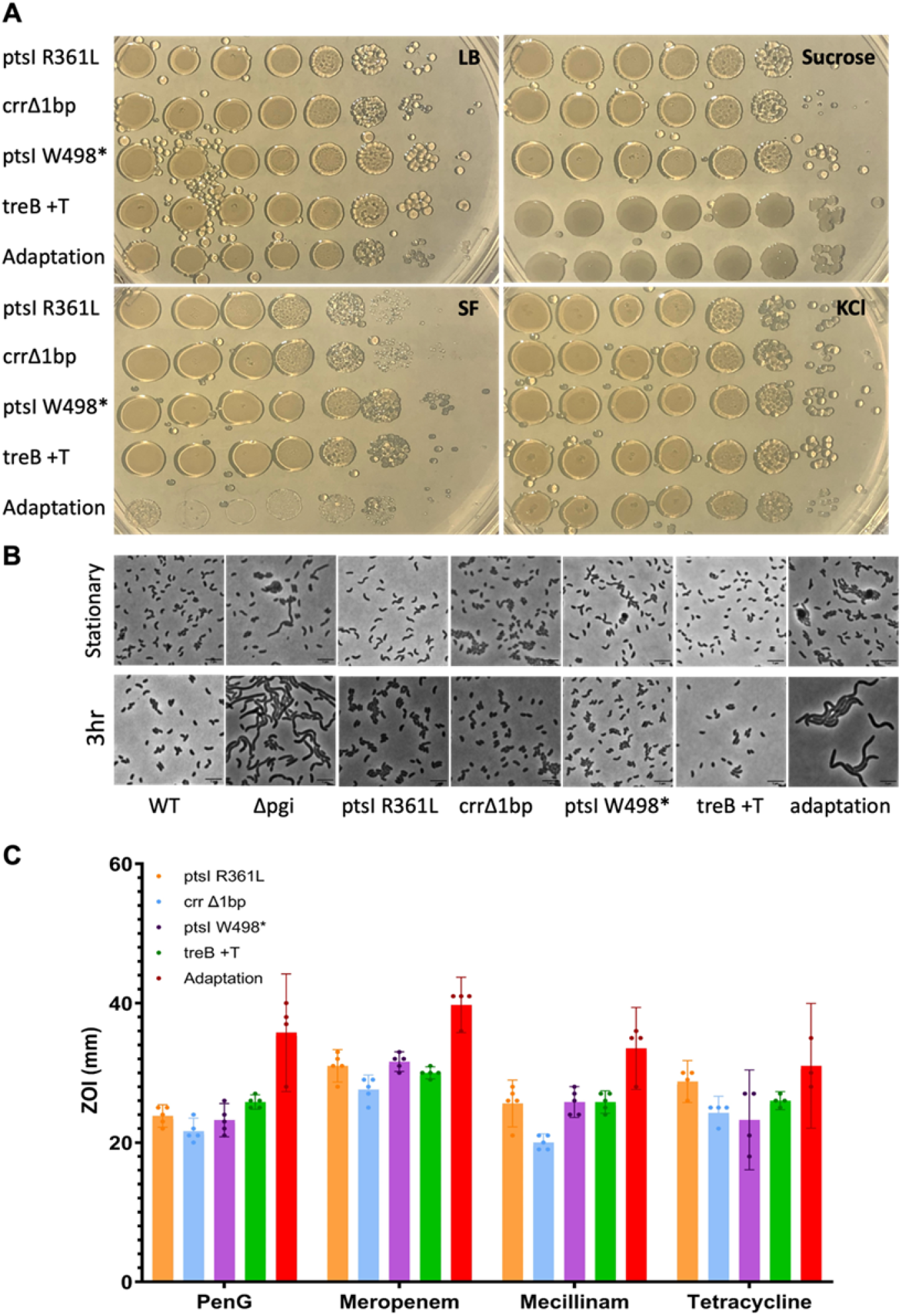
Spontaneous suppressors in PTS restore *pgi* defects. A) Overnight cultures of the indicated strains were serially diluted and spotted on LB, SF, SF + 180mM sucrose, and SF + 200mM KCl B) The indicated strains were diluted 100fold into fresh LB, then imaged on an agarose pad after 3 hours of growth. Scale bar = 5μm C) Zone of inhibition measurements from a disk diffusion experiment.

**Supplemental Figure 3:**
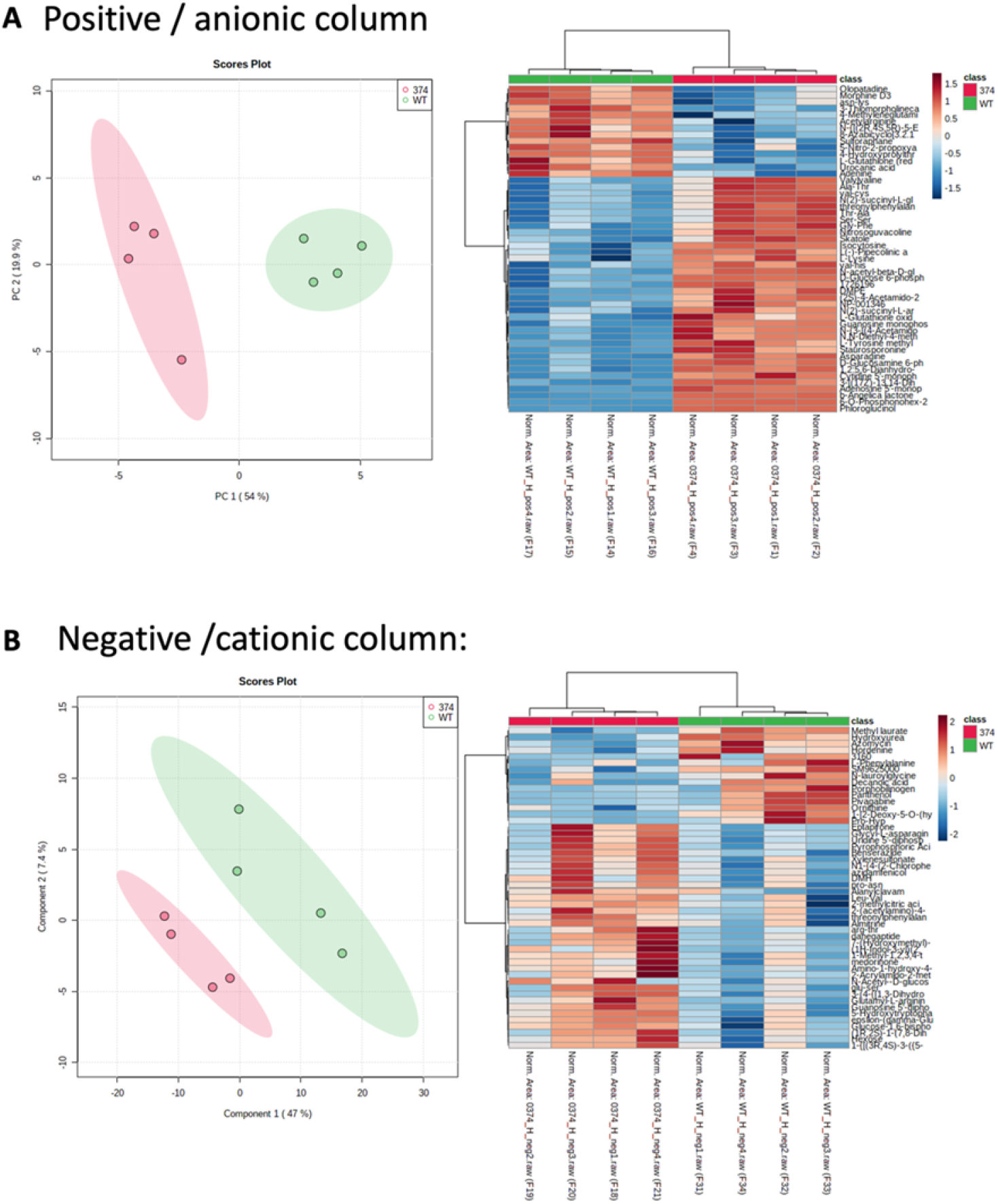
Top metabolomic variations between WT and *pgi*. Heatmap and QC plot of metabolite differences between WT and *pgi* on positive and negative HILIC columns. n = 4.

**Supplemental Figure 4:**
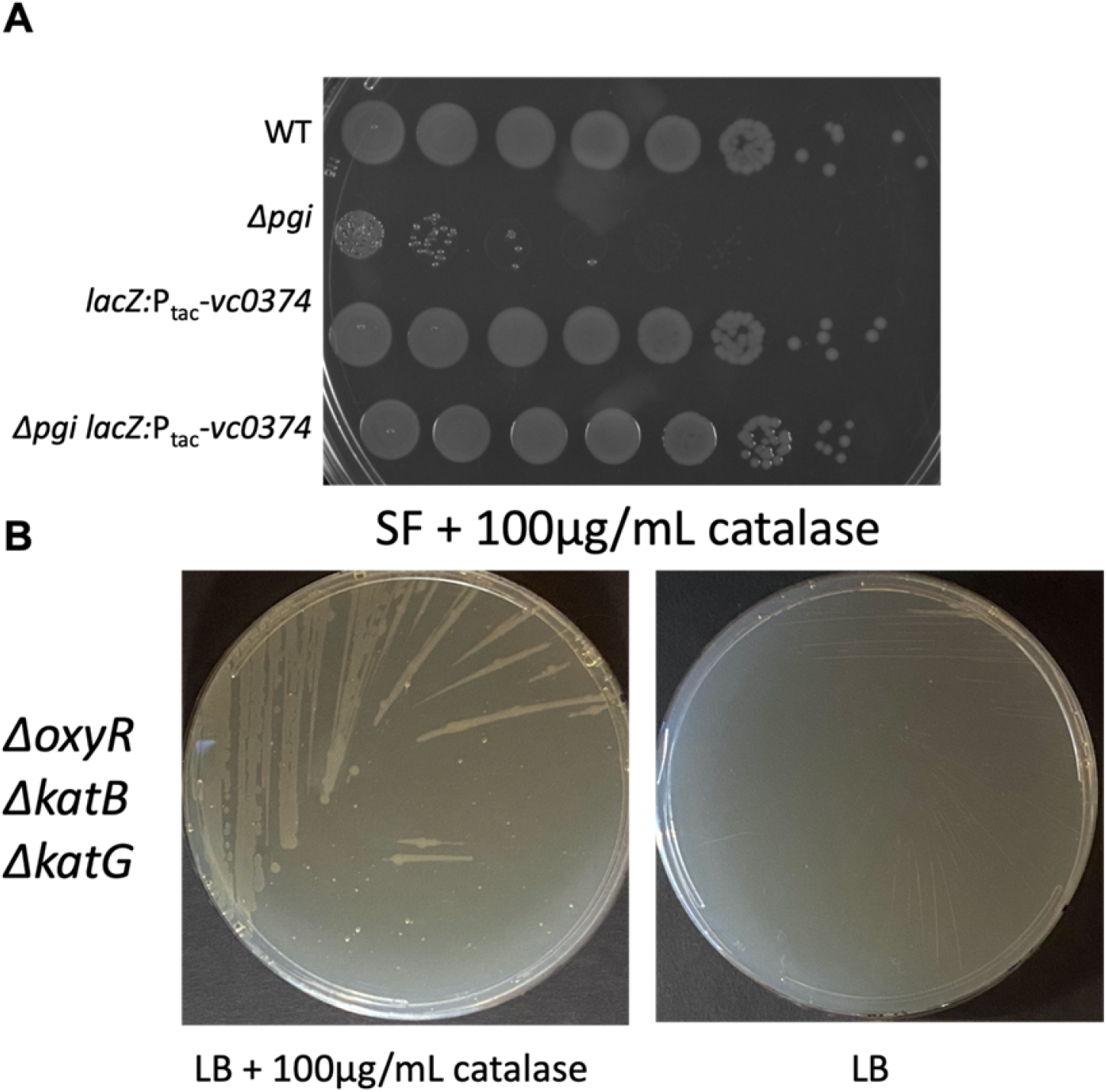
*pgi* defects are not due to excessive ROS. Serial dilutions of the indicated strains were spotted on a SF plate containing 100μg/mL catalase to degrade reactive oxygen species. A highly ROS-sensitive strain (Δ*oxyR* Δ*katB* Δ*katG*) was applied to a plate from the same batch as a control for functional catalase in the plate.

**Supplemental Figure 5:**
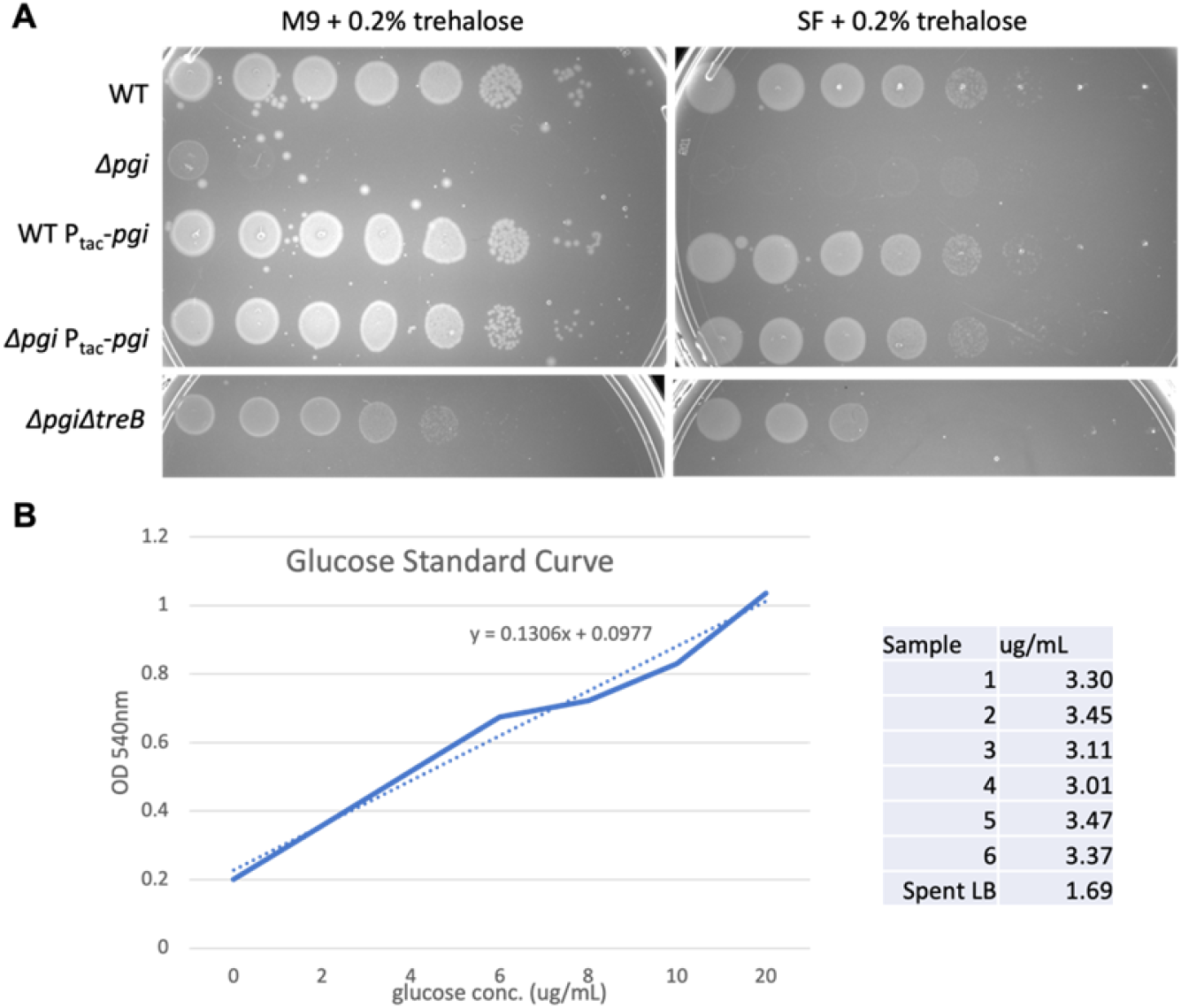
Exploring the effects of trehalose and glucose in LB and their effect on *pgi* mutants. A). Serial dilutions of the indicated strains were plated on SF or M9 containing 0.2% trehalose. B) Glucose was quantified using GO Assay kit (See Methods). Shown are glucose concentrations (μg/mL) of 6 different batches of LB broth and the respective standard curve.

**Supplemental Table 1: Summary of genes that answered the vxrAB induction screen**

Shown are all transposon disruptions that answered the screen, organized by reporter strain background (P_*vca0140*_-*lacZ* or P_*vxrA*_-*lacZ*), predicted gene function, and the frequency of occurrence.

**Supplemental Table 2: Spontaneous Suppressor mutations reveal PTS target**

Suppressor mutations were identified by whole genome sequencing. Shown are suppressors with their respective mutations and the frequency in which they occurred.

## References

1. Brauner A, Fridman O, Gefen O, Balaban NQ. 2016. Distinguishing between resistance, tolerance and persistence to antibiotic treatment. Nat Rev Microbiol 14:320–330.

2. Lazarovits G, Gefen O, Cahanian N, Adler K, Fluss R, Levin-Reisman I, Ronin I, Motro Y, Moran-Gilad J, Balaban NQ, Strahilevitz J. 2022. Prevalence of Antibiotic Tolerance and Risk for Reinfection Among E. coli Bloodstream Isolates: A Prospective Cohort Study. Clin Infect Dis Off Publ Infect Dis Soc Am 75(10):1706–1713.

3. Levin-Reisman I, Ronin I, Gefen O, Braniss I, Shoresh N, Balaban NQ. 2017. Antibiotic tolerance facilitates the evolution of resistance. Science 355:826–830.

4. Sulaiman JE, Lam H. 2021. Evolution of Bacterial Tolerance Under Antibiotic Treatment and Its Implications on the Development of Resistance. Front Microbiol 12.

5. Windels EM, Michiels JE, Van den Bergh B, Fauvart M, Michiels J. 2019. Antibiotics: Combatting Tolerance To Stop Resistance. mBio 10:e02095–19.

6. Reygaert WC. 2018. An overview of the antimicrobial resistance mechanisms of bacteria. AIMS Microbiol 4:482–501.

7. Cross T, Ransegnola B, Shin J-H, Weaver A, Fauntleroy K, VanNieuwenhze MS, Westblade LF, Dörr T. 2019. Spheroplast-Mediated Carbapenem Tolerance in Gram-Negative Pathogens. Antimicrob Agents Chemother 63:e00756–19.

8. Dörr T. Understanding tolerance to cell wall–active antibiotics. 2021. Ann NY Acad Sci. 1496:35–58.

9. Typas A, Banzhaf M, Gross CA, Vollmer W. 2011. From the regulation of peptidoglycan synthesis to bacterial growth and morphology. Nat Rev Microbiol 10:123–136.

10. Weaver AI, Murphy SG, Umans BD, Tallavajhala S, Onyekwere I, Wittels S, Shin J-H, VanNieuwenhze M, Waldor MK, Dörr T. 2018. Genetic Determinants of Penicillin Tolerance in Vibrio cholerae. Antimicrob Agents Chemother 62.

11. Dörr T, Davis BM, Waldor MK. 2015. Endopeptidase-Mediated Beta Lactam Tolerance. PLOS Pathog 11:e1004850.

12. Monahan LG, Turnbull L, Osvath SR, Birch D, Charles IG, Whitchurch CB. 2014. Rapid conversion of Pseudomonas aeruginosa to a spherical cell morphotype facilitates tolerance to carbapenems and penicillins but increases susceptibility to antimicrobial peptides. Antimicrob Agents Chemother 58:1956–1962.

13. Cheng AT, Ottemann KM, Yildiz FH. 2015. Vibrio cholerae Response Regulator VxrB Controls Colonization and Regulates the Type VI Secretion System. PLOS Pathog 11:e1004933.

14. Dörr T, Alvarez L, Delgado F, Davis BM, Cava F, Waldor MK. 2016. A cell wall damage response mediated by a sensor kinase/response regulator pair enables beta-lactam tolerance. Proc Natl Acad Sci 113:404–409.

15. Shin J-H, Choe D, Ransegnola B, Hong H-R, Onyekwere I, Cross T, Shi Q, Cho B-K, Westblade LF, Brito IL, Dörr T. 2021. A multifaceted cellular damage repair and prevention pathway promotes high level tolerance to β-lactam antibiotics. EMBO Rep 22(2):e51790.

16. Peschek N, Herzog R, Singh PK, Sprenger M, Meyer F, Fröhlich KS, Schröger L, Bramkamp M, Drescher K, Papenfort K. 2020. RNA-mediated control of cell shape modulates antibiotic resistance in Vibrio cholerae. Nat Commun 11:6067.

17. Teschler JK, Cheng AT, Yildiz FH. 2017. The Two-Component Signal Transduction System VxrAB Positively Regulates Vibrio cholerae Biofilm Formation. J Bacteriol 199.

18. Cho SY, Yoon S. 2020. Structural analysis of the sensor domain of the β-lactam antibiotic receptor VbrK from Vibrio parahaemolyticus. Biochem Biophys Res Commun 533:155– 161.

19. Li L, Wang Q, Zhang H, Yang M, Khan MI, Zhou X. 2016. Sensor histidine kinase is a β-lactam receptor and induces resistance to β-lactam antibiotics. Proc Natl Acad Sci U S A 113:1648–1653.

20. Goh BC, Chua YK, Qian X, Lin J, Savko M, Dedon PC, Lescar J. 2020. Crystal structure of the periplasmic sensor domain of histidine kinase VbrK suggests indirect sensing of β-lactam antibiotics. J Struct Biol 212:107610.

21. Tan K, Teschler JK, Wu R, Jedrzejczak RP, Zhou M, Shuvalova LA, Endres MJ, Welk LF, Kwon K, Anderson WF, Satchell KJF, Yildiz FH, Joachimiak A. Sensor Domain of Histidine Kinase VxrA of Vibrio cholerae: Hairpin-Swapped Dimer and Its Conformational Change. J Bacteriol 203:e00643–20.

22. Harper CE, Zhang W, Shin J-H, Wijngaarden E van, Chou E, Lee J, Wang Z, Dörr T, Chen P, Hernandez CJ. 2022. Mechanical stimuli activate gene expression via a cell envelope stress sensing pathway. bioRxiv https://doi.org/10.1101/2022.09.25.509347.

23. Hernández SB, Dörr T, Waldor MK, Cava F. 2020. Modulation of Peptidoglycan Synthesis by Recycled Cell Wall Tetrapeptides. Cell Rep 31:107578.

24. Sit B, Srisuknimit V, Bueno E, Zingl FG, Hullahalli K, Cava F, Waldor MK. 2022. Undecaprenyl phosphate translocases confer conditional microbial fitness. Nature 1–3. https://doi.org/10.1038/s41586-022-05569-1

25. Dörr T, Lam H, Alvarez L, Cava F, Davis B, Waldor M. 2014. A Novel Peptidoglycan Binding Protein Crucial for PBP1A-Mediated Cell Wall Biogenesis in Vibrio cholerae. PLoS Genet 10:e1004433.

26. Caparrós M, Pisabarro AG, de Pedro MA. 1992. Effect of D-amino acids on structure and synthesis of peptidoglycan in Escherichia coli. J Bacteriol 174:5549–5559.

27. Ho BT, Basler M, Mekalanos JJ. 2013. Type 6 Secretion System–Mediated Immunity to Type 4 Secretion System–Mediated Gene Transfer. Science 342:250–253.

28. Hayes CA, Dalia TN, Dalia AB. 2017. Systematic genetic dissection of PTS in Vibrio cholerae uncovers a novel glucose transporter and a limited role for PTS during infection of a mammalian host. Mol Microbiol 104:568–579.

29. Boulanger EF, Sabag-Daigle A, Thirugnanasambantham P, Gopalan V, Ahmer BMM. 2021. Sugar-Phosphate Toxicities. Microbiol Mol Biol Rev 85:e00123–21.

30. Achkar J, Xian M, Zhao H, Frost JW. 2005. Biosynthesis of Phloroglucinol. J Am Chem Soc 127:5332–5333.

31. Christodoulou D, Link H, Fuhrer T, Kochanowski K, Gerosa L, Sauer U. 2018. Reserve Flux Capacity in the Pentose Phosphate Pathway Enables Escherichia coli’s Rapid Response to Oxidative Stress. Cell Syst 6:569–578.

32. Sezonov G, Joseleau-Petit D, D’Ari R. 2007. Escherichia coli physiology in Luria-Bertani broth. J Bacteriol 189:8746–8749.

33. Rimmele M, Boos W. 1994. Trehalose-6-phosphate hydrolase of Escherichia coli. J Bacteriol 176:5654–5664.

34. Zhou Y, Yan K, Qin Q, Raimi OG, D. C, Wang B, Ahamefule CS, Kowalski B, Jin C, van Aalten DMF, Fang W. 2022. Phosphoglucose Isomerase Is Important for Aspergillus fumigatus Cell Wall Biogenesis. mBio 13:e01426–22.

35. Tuckman D, Donnelly RJ, Zhao FX, Jacobs WR, Connell ND. 1997. Interruption of the phosphoglucose isomerase gene results in glucose auxotrophy in Mycobacterium smegmatis. J Bacteriol 179:2724–2730.

36. Prasad C, Freese E. 1974. Cell Lysis of Bacillus subtilis Caused by Intracellular Accumulation of Glucose-1-Phosphate. J Bacteriol 118:1111–1122.

37. Yadav V, Panilaitis B, Shi H, Numuta K, Lee K, Kaplan DL. 2011. N-acetylglucosamine 6-Phosphate Deacetylase (nagA) Is Required for N-acetyl Glucosamine Assimilation in Gluconacetobacter xylinus. PLOS ONE 6:e18099.

38. Meibom KL, Li XB, Nielsen AT, Wu C-Y, Roseman S, Schoolnik GK. 2004. The Vibrio cholerae chitin utilization program. Proc Natl Acad Sci 101:2524–2529.

39. Sloan R, Surber J, Roy EJ, Hartig E, Morgenstein RM. 2022. Enzyme 1 of the phosphoenolpyruvate:sugar phosphotransferase system is involved in resistance to MreB disruption in wild-type and ΔenvC cells. Mol Microbiol 118:588–600.

40. Sachla AJ, Helmann JD. 2019. A bacterial checkpoint protein for ribosome assembly moonlights as an essential metabolite-proofreading enzyme. Nat Commun 10:1526.

41. Espinosa E, Daniel S, Hernández SB, Cava F, Barre F-X, Galli E. 2020. L-arabinose induces the formation of viable non-proliferating Vibrio cholerae spheroplasts. bioRxiv 2020.07.15.203711.

42. Stokes JM, Lopatkin AJ, Lobritz MA, Collins JJ. 2019. Bacterial Metabolism and Antibiotic Efficacy. Cell Metab 30:251–259.

43. Ferrières L, Hémery G, Nham T, Guérout A-M, Mazel D, Beloin C, Ghigo J-M. 2010. Silent Mischief: Bacteriophage Mu Insertions Contaminate Products of Escherichia coli Random Mutagenesis Performed Using Suicidal Transposon Delivery Plasmids Mobilized by Broad-Host-Range RP4 Conjugative Machinery. J Bacteriol 192:6418–6427.

44. Gibson DG, Young L, Chuang R-Y, Venter JC, Hutchison CA, Smith HO. 2009. Enzymatic assembly of DNA molecules up to several hundred kilobases. Nat Methods 6:343–345.

45. Lazarus JE, Warr AR, Kuehl CJ, Giorgio RT, Davis BM, Waldor MK. 2019. A New Suite of Allelic-Exchange Vectors for the Scarless Modification of Proteobacterial Genomes. Appl Environ Microbiol 85:e00990–19.

46. Miyata ST, Unterweger D, Rudko SP, Pukatzki S. 2013. Dual Expression Profile of Type VI Secretion System Immunity Genes Protects Pandemic Vibrio cholerae. PLOS Pathog 9:e1003752.

47. Wilson AC, Perego M, Hoch JA. 2007. New transposon delivery plasmids for insertional mutagenesis in Bacillus anthracis. J Microbiol Methods 71:332–335.

